# Intramolecular synergism and bacterial lysis regulation can explain the complex multi-domain architecture of the bacteriophage endolysin PlySK1249

**DOI:** 10.1101/2020.12.01.406769

**Authors:** Frank Oechslin, Carmen Menzi, Philippe Moreillon, Gregory Resch

**Affiliations:** Dept. of Fundamental Microbiology, University of Lausanne, Switzerland; Dept. of Biochemistry, Microbiology and Bioinformatics, Université Laval, Canada

**Author notes:** Corresponding author: Frank Oechslin.

**Keywords:** Bacteriophage, endolysin, PlySK1249, intramolecular synergism, lysis regulation, proteolysis

## Abstract

Endolysins are peptidoglycan hydrolases produced at the end of the bacteriophage (phage) replication cycle to lyse the host cell. Gram-positive phages endolysins come in a variety of multi-modular forms that combine different catalytic domains and may have evolved to adapt to their bacterial hosts. However, the reason why phage can adopt endolysin with such complex multidomain architecture is for the moment not well understood.

We used the *Streptococcus dysgalactiae* phage endolysin PlySK1249 as a model to study the implication of multi-domain architecture in phage-induced bacterial lysis and lysis regulation. The activity of the enzyme relied on a bacteriolytic amidase (Ami), a non-bacteriolytic L-Ala-D-Ala endopeptidase (CHAP) acting as a de-chaining enzyme and central LysM cell wall binding domain (CBD).

Ami and CHAP synergized for peptidoglycan digestion and bacteriolysis in the native enzyme or when expressed individually and reunified *in vitro*. This cooperation could be modulated by bacterial cell wall-associated proteases, which specifically cleaved the two linkers connecting the different domains. While both catalytic domains were observed to act coordinately to optimize bacterial lysis, the CBD is expected to delay diffusion of the enzyme until proteolytic inactivation is achieved.

As for certain autolysins, PlySK1249 cleavage by bacterial cell wall associated proteases might be an example of dual phage-bacterial regulation and mutual coevolution. In addition, understanding more thoroughly the multidomain interplay of PlySK1249 opens new perspectives on the ideal architecture of therapeutic antibacterial endolysins.

## Introduction

Phage endolysins represent a fascinating family of peptidoglycan hydrolases that are critical for bacterial lysis and release of phage progeny at the end of the phage life cycle (1). They have gained recent interest in biomedical development as potential antibacterial agents (2–4). Phage endolysins target the bacterial peptidoglycan, an essential structure made of a complex meshwork of *N*-acetylglucosamine (GlcNAc)–*N*-acetylmuramic acid (MurNAc) glycan strands cross-linked by short stem peptides attached to MurNAc residues (5,6). They can cleave various bonds between sugars or peptide chains and come in a variety of uni- or multi-modular forms that may have evolved to adapt to their bacterial hosts. Endolysins encoded by phages infecting Gram-positive bacteria are usually composed of several functional domains including glucosaminidases, amidases and endopeptidases (referred to as catalytic domains or CDs) and cell wall-binding domains (referred to as CBDs)(7–9). However, the architecture and functions of CDs and CBDs are quite variable. For instance, the well-characterized staphylococcal endolysins LysK, Ф11, and MV-L, as well as the streptococcal endolysins B30, PlyGBS and lambda Sa2 contain two CDs (10–15) in spite only one induces bacterial lysis of cell cultures (7,16–19).

The reasons for conserving non-bacteriolytic CDs are not entirely clear. It has been hypothesized that their lytic activities were progressively lost through evolution, or that they might be involved in reinforcing cell wall binding (15,20). In symmetry, CBDs – which are not bacteriolytic by themselves – are thought to be important for addressing and positioning the CDs in the peptidoglycan and conferring the endolysin specificity to their target bacteria (21–24). They are commonly located at the N- or C-terminus of the protein and are rarely located in the central region (25,26). However, CBDs are not always mandatory for bacterial lysis since deleting them has even improved the lytic activity of some endolysins even without loss of specificity, as shown for endolysin B30, (7,17,18). Hence, non-bacteriolytic CDs or CBDs may have additional or alternative roles in the endolysin physiology.

In this work, we attempted to clarify the functional roles of the CDs and the CBD of the *Streptococcus dysgalactiae* phage endolysin PlySK1249 (27) and their implication in the physiology of phage-induced bacterial lysis. This endolysin has a complex structure with a central LysM CBD that is bracketed by a N-terminal amidase (Ami) domain and a C-terminal Cysteine Histidine-dependent Amidohydrolase Peptidase (CHAP) domain. While amidases (or *N*-acetylmuramoyl-L-alanine amidases) are enzymes known to hydrolyze the amide bond between the *N*-acetylmuramoyl glycan moiety and the first L-ala of the stem peptides (28), CHAP domains are primarily endopeptidases, sometimes with amidase activities (29).

We could observe that the Ami domain was a bacteriolytic amidase whereas CHAP was a non-bacteriolytic L-Ala-D-Ala endopeptidase that could synergize for efficient bacterial lysis. The CHAP domain also acted as a de-chaining enzyme and shared high functional and structural analogies with de-chaining autolysins present in lactococci or streptococci (30,31). We finally notice that PlySK1249 was subject to proteolytic cleavage by cell wall proteases both *ex vivo* and after phage induction *in vivo*. Cleavage dismantled the CDs by hydrolyzing their linker regions, thus hindering their bacteriolytic cooperation and possibly modulate the lytic activity of the enzyme.

Like for proteolytic regulation of certain autolysins (32–34), PlySK1249 cleavage could well represent a new mechanism of dual phage-bacterial regulation of endolysin-induced lysis. In addition, understanding more thoroughly the inter-domain interplay of PlySK1249 may be useful in the design of novel therapeutic lysins.

## Results

### Contribution and cooperation of each domains in the activity of the PlySK1249 endolysin

To assess the specific roles of each of the predicted domains, various truncated forms of PlySK1249 were generated (Fig. 1A). After being overexpressed in *Escherichia coli*, the protein constructs could be purified by affinity chromatography using a 6X-His Tag at the C-terminal position. The purity and the correct molecular weight were verified on 4-12% BisTris gels (Fig. 1B) (~54 kDa for PlySK1249, ~19 kDa for Ami, ~37 kDa for Ami_LysM, ~26 kDa for LysM_CHAP). Of note, the CHAP domain could not be overexpressed alone, it always required attachment to the LysM domain.

**Figure 1.**
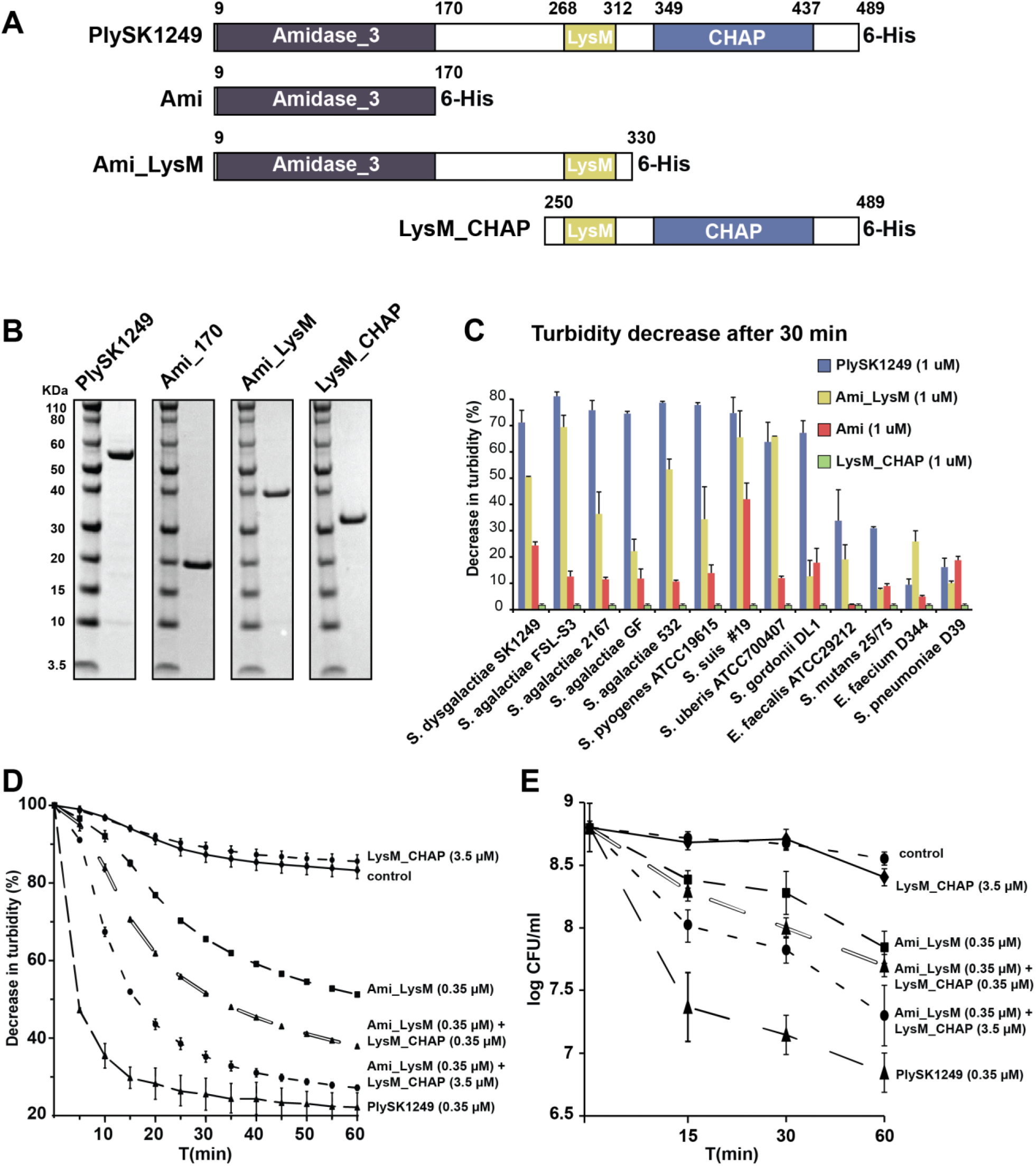
Characterization of the lytic activity of the PlySK1249 endolysin and its various truncated forms. A) Parent PlySK1249 (full enzyme, aa 1-489); Ami (N-terminal amidase, aa 1-170); Ami_LysM (N-terminal amidase + LysM, aa 1-330); LysM_CHAP (LysM + C-terminal CHAP domain, aa 250-489). B) SDS-PAGE gel showing PlySK1249 and its various truncated constructions. The purified endolysin constructions were loaded (2 mg/ml) on NuPAGE 4-12% BisTris gels and stained with Coomassie blue. Molecular mass was determined with a pre-stained protein standard. C) Cells of multiple bacterial species in the exponential growth phase were exposed to 1 μM of parent PlySK149 or its different truncated forms. Decreases in the turbidity of the cultures were measured at 600 nm after 30 min. No decrease in turbidity was observed in the absence of enzyme (not shown) or in the presence of LysM_CHAP. D) *S. dysgalactiae* cells in the exponential growth phase were also exposed to either 1 μM of Ami_LysM or LysM_CHAP constructs alone or in combination and the decreases in turbidity (at 600 nm) was followed over 1 h. E) Viable counts from experiment shown in panel D). Counts were assessed by plating serial dilutions on nutrient agar. The experiments were repeated twice in triplicate and means ± standard deviations are shown.

The relative contribution of the two CDs and the CBD to bacterial lysis was compared to the relative contribution of the parent enzyme. The constructs were first tested against an array of different Gram-positive bacteria (Figure 1C). Parent PlySK1249 was more lytic against *Streptococcus* spp. than against more distantly related species, such as enterococci. Moreover, and apart from a few exceptions, the PlySK1249 truncated forms were less active than the parent enzyme, with Ami_LysM being generally more active than Ami alone and LysM-CHAP showing no lytic activity even when incubated for 2 h (Fig. 1C and data not shown).

We further tested the possibility of a cooperative activity between the Ami and CHAP catalytic domains by following the loss of turbidity and cell viability when used alone or mixed together against *S. dysgalactiae* 1249 cells (Fig. 1D and 1E, respectively). The results confirmed the decrease of intrinsic lytic activity of the subdomains observed in Fig. 1C (Fig. 1D). Indeed, while PlySK1249 caused a rapid 50% drop in turbidity within 5 min, Ami_LysM showed intermediate lysis (50% drop in turbidity in 1 h) and LysM_CHAP was not lytic at all. However, adding equimolar amounts of non-lytic LysM_CHAP to Ami_LysM gradually increased lysis while a 1/10 Ami_LysM/LysM_CHAP ratio almost completely restoring lysis efficiency at 1 h, as compared to the parent PlySK1249. This inter-domain cooperation was formally synergistic (i.e. the effect of the domain combination was superior than the sum of their individual effects) and was also observed between Ami and LysM-CHAP (data not presented). These results correlated with the viable cell counts obtained in time-kill assays (Fig. 1E). Thus, although LysM_CHAP had no intrinsic lytic activity, it clearly contributed to the overall activity of the PlySk1249 endolysin.

### The non-lytic CHAP domain has an intrinsic de-chaining activity

A closer look at the LysM_CHAP activity using optical and Transmission Electron Microscopy (TEM) revealed subtle non-lytic effects (Fig. 2). Phase-contrast optical microscopy of exponential phase cultures (OD_600nm_ 0.5) of *S. dysgalactiae* SK1249 showed that they were composed of 90% of chains ranging from of 6 to 37 cells (Fig. 2A.I and Fig. 2A.II). However, 1 h of incubation with LysM-CHAP resulted in chain disruption with the majority (60%) of chains harboring only 1 to 5 bacteria, the rest of them being distributed over longer structures (Fig. 2A.III and Fig. 2A.IV).

**Figure 2.**
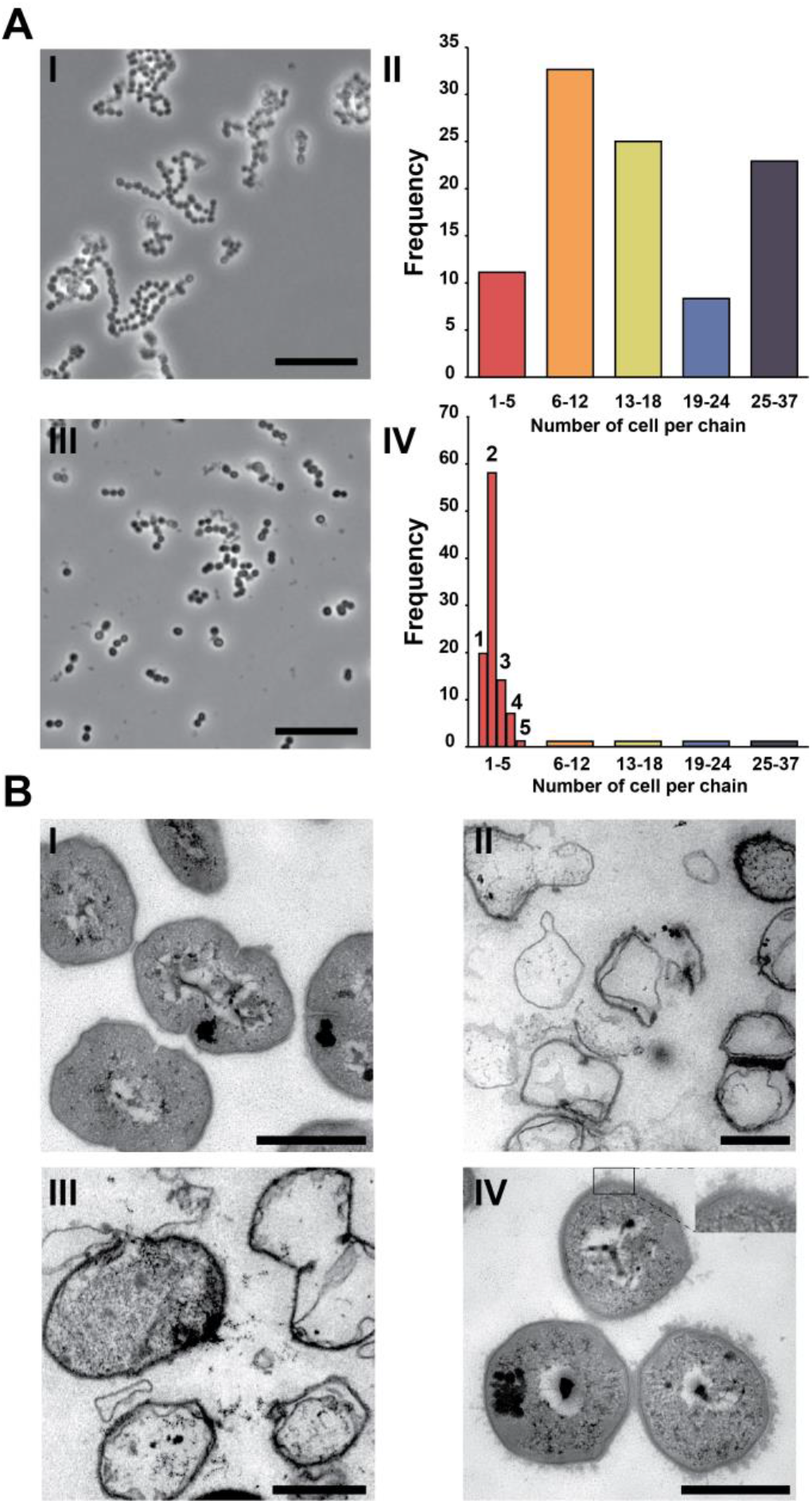
Morphological aspects of cells lysed by the PlySK1249 endolysin and its various truncated forms. A) *S. dysgalactiae* in the exponential growth phase were observed after one hour using phosphate buffer as a control (A.I), or LysM_CHAP at a final concentration of 3.5 μM (A.III). Although the CHAP domain did not impact directly on cell lysis, an effect on chain disruption was observed. The control population was composed mainly (90%) of chains between 6 and 37 cells long (144 cells observed in total, A.II) compared to the 60% of chains that were composed of two cells for the LysM_CHAP treatment (1044 cells observed in total, A.IV). B) Transmission electron microscopy of *S. dysgalactiae* cells treated with the native PlySK1249 endolysin or its various truncated versions. *S. dysgalactiae* cells in the exponential growth phase were treated with (B.I) phosphate buffer for 1 h as a control, (B.II) Plysk1249 for 15 min, (B.III) Ami_LysM for 15 min, or (B.IV) LysM_CHAP for 1 h. Cells were then post-fixed using glutaraldehyde and embedded in epoxy for ultrathin sectioning. Scale bars represent 500 nm. The inset in figure B.IV highlights the non-lytic wall nibbling by the CHAP endopeptidase.

This non-lytic activity was further supported by TEM imaging. Control cells incubated in PBS alone showed a typical Gram-positive bacterial shape (Fig. 2B.I). Cells treated with PlySK1249 or Ami_LysM were lysed as assessed by the presence of a majority of ghost cells (15 min incubation, Fig. 2B.II and 2B.III, respectively). Treatment with LysM_CHAP alone resulted in round swelling cells with surface alterations but absence of lysis even after 1 h exposure (Fig. 2B.IV).

### The N-terminal Ami domain of PlySK1249 is an amidase while the C-terminal CHAP domain is an endopeptidase

To better understand to contribution of each CD domains to the overall activity of the enzyme, we further looked at their specific peptidoglycan cleavage sites. Glycosidase or peptidase activity was first assed for PlySK1249 and its truncated constructs using purified *S. dysgalactiae* cell wall. Compared to the N-acetylmuramidase mutanolysin, none of the constructs had glycosidase activities according to Park-Johnson assay (Supplementary Table 2). In contrast, using a modified Ghuysen assay, free amino groups release was measured after peptidoglycan hydrolysis by PlyKS1249 and all the PlySK1249-derived constructs, therefore identifying an amidase and/or endopeptidase activity.

To discriminate between amidase and endopeptidase activities for the Ami and CHAP catalytic domains, digested peptidoglycan was further analyzed RP-HPLC and LC-MS for the presence of precursor masses containing the stem peptide motif AQKAAA and polymers of it. After digestion with PlySK1249, two major peaks with relatively short retention times were identified by RP-HPLC (at approximately 35 and 42 min, Fig. 3A.I). These were mainly identified as stem peptides dimers by LC-MS also presence of monomers and trimers was detected (Fig. 3.B). After digestion with Ami_LysM or Ami alone, the two major peaks became marginal and were replaced by an array of smaller peaks eluting later in the RF-HPLC profile (between 55 to 90 min, Fig. 3A.II and Supplementary Fig. 1, respectively). LC-MS analysis showed that these peaks covered an array from trimers to heptamers that eluted later in RF-HPLC profiles (Fig. 3B and Supplementary Fig. 2). Moreover, identical profiles were obtained for Ami and Ami_LysM, indicating that the LysM CBD was not responsible for altering the hydrolytic profile.

**Figure 3.**
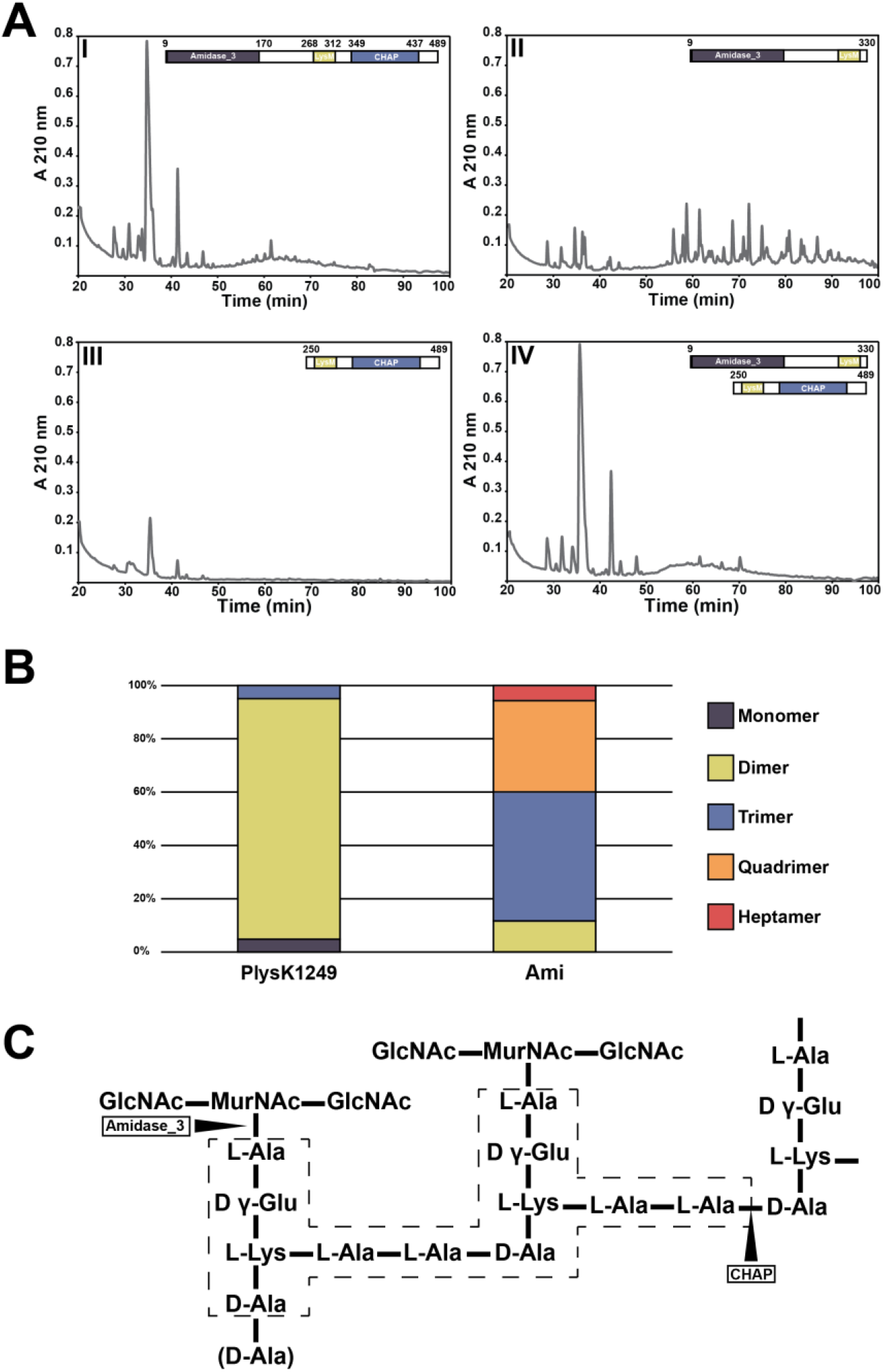
RP-HPLC chromatogram and LC-MS analysis of *S. dysgalactiae* peptidoglycan digested with PlySK1249 or its truncated catalytic domains. A) Purified wall-peptidoglycan was digested overnight and glycans were sequentially precipitated before chromatography on a C18 Sephasil column. Equimolar concentrations (3.5 μM) of I) PlySK1249, II) Ami_LysM, III) LysM_CHAP and IV) both Ami_LysM and LysM_CHAP were used and analyses were repeated three times for each, yielding the same results. B) Relative abundances of the polymers observed in the native enzyme and amidase domain digestion experiments. The products of peptidoglycan digestion by the native enzyme and its truncated Ami domain were analysed by LC-MS after a 5 kDa filtration. The masses of the precursors corresponding to the dimer, trimer, quadrimer and heptamer of the AAAQKA monomer block were detected. All of these oligomers were detected in both samples, with the exception of the heptamers, which were only present in the amidase digestion. Polymer structures were deduced from masses after *de novo* peptide sequencing. C) Representation of *S. dysgalactiae* peptidoglycan based on the masses and sequences observed. Arrows indicate the cleavage sites for the respective catalytic domains of the enzyme. The framed dotted-line area corresponds to the dimer block

Importantly, while digestion with LysM_CHAP did not yield any major peak pattern (Fig. 3A.III), combining LysM_CHAP with Ami-LysM restored the complete digestion pattern of the whole PlySK1249 enzyme (Fig. 3A.IV). This observation indicates that LysM_CHAP could resolve stem-peptide multimers mainly to dimers, but hardly to monomers. Since glycan strands can remain connected through dimer crosslinks, which are sufficient to maintain a loose polymer structure and thus prevent bacterial lysis, this might explain why the CHAP domain is not directly lytic on its own. Finally, the digestion products summarized in Supplementary Fig. 2 indicate an N-acetylmuramoyl-L-alanine activity for the Ami domain and a L-Ala-D-Ala endopeptidase activity for the CHAP domain (Fig. 3C).

### PlySK1249 is composed of proteolytic-resistant core domains connected by proteolytic-susceptible linkers

The complex nature of PlySK1249 may be a source of instability as previous authors observed degradation of multi-domain autolysins by cell-wall associated proteases (32–34). For this reason, we tested the resistance of PlySK1249 to protease-induced cleavage first by trypsin and second by bacterial wall associated proteases.

Proteolytic cleavage of PlySK1249 was observed to be restricted to the two linkers connecting the different enzyme active domains, while the active domains themselves were left intact. Two bands having a lower molecular mass could be observed after overnight incubation of the endolysin with uM/ml of trypsin (Fig. 4A). Amino acid sequencing of extracted bands confirmed the presence of two peptides corresponding to the CHAP (aa 318 to 489) and Ami (aa 1 to 204) CD domains in the upper band and the Ami (aa 1 to 175) domain only in the lower band. These results were also confirmed by western blotting using Anti-6X His antibodies. Similar results were finally observed when the Ami_LysM or the LysM_CHAP constructs were exposed to trypsin degradation, which also resulted with the cleavage of the connecting linkers (Fig. 4B and 4C). Thus, PlySK1249 was composed of different trypsin-resistant cores corresponding to the CD and CBD domains, and trypsin-susceptible linker regions.

**Figure 4.**
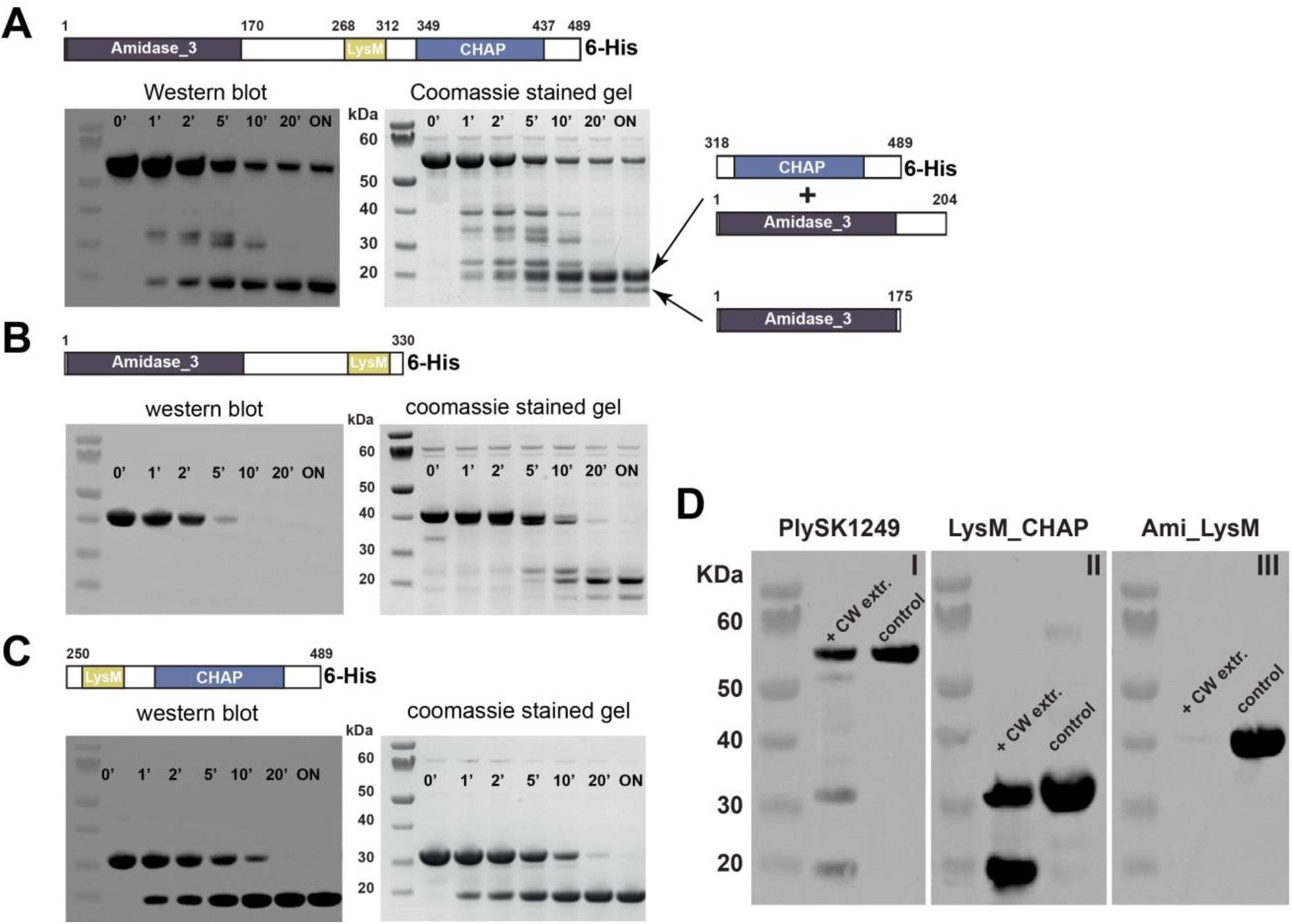
PlySK1259 proteolysis and its different truncated versions in the presence of trypsin or cell wall associated proteases. 40 μM of the parent enzyme (A) or the Ami_lysM (B) and LysM_CHAP (C) were incubated with 0.1 uM/ml of trypsin. Right panels: Sample were migrated after different incubation times on NuPAGE 4-12% BisTris, stained with Coomassie blue and subsequent degradation products were sequenced. Left panels: digestion products were transferred to a nitrocellulose membrane for western blotting using Anti-6X His antibody. D) 40 μM of the parent enzyme (I), Ami_lysM (II) or LysM_CHAP (III) were incubated overnight with cell wall protein extract from *S. dysgalactiae* SK1249. Samples were migrated on NuPAGE 4-12% BisTris and degradation was confirmed by western blotting using Anti-6X His tag antibody.

Extracts of *S. dysgalactiae* SK1249 cell wall-associated proteins were also prepared and incubated with either the native enzyme or the Ami-LysM and LysM-CHAP constructs (Fig. 4D). Due to the large number of proteins in the cell wall extract, it was not possible to visualize the degradation products by Coomassie-blue staining. Therefore, Western blotting with an Anti-6X His tag antibody was used as for trypsin digestion. After overnight incubation, two bands were generated for PlySK1249 at approximately 30 and 20 kDa, which were absent in the control with the endolysin alone, as well as in the wall protein extract (result not shown) (Fig. 4D.I). Degradation was also observed for the LysM-CHAP construct with an additional band at 20 kDa (Fig. 4D.II). Due to the position of the His-Tag in the C-terminal part of the enzyme, the Ami domain alone was not observed (Fig. 4D.III). Importantly, these results are similar to the one observed using trypsin and confirmed that PlySK1249 was cleaved between its two linkers, both by trypsin and cell-wall proteases.

### Insights into the nature of the wall proteases

We used a bioactivity-guided fractionation protocol to identify the host cell wall associated proteases that are responsible for PlySK1249 degradation (Fig. 5. Ammonium sulfate was used for a first round of sorting and fractions were tested on the LysM_CHAP construct. LysM_CHAP proteolytic degradation was observed for fractions precipitated at 60% to 75% sulfate ammonium (Fig. 5A). A second purification step was performed by size exclusion chromatography using a 55% to 80% ammonium sulfate fraction and the output was again tested for proteolytic activity on the LysM_CHAP construct (Fig. 5B). A total of four fractions were analyzed by LC-MS. These were fraction B4 and B6, in which no activity was observed, and fractions B8 and B10, in which proteolytic activity was observed (Fig. 5C). After LC-MS analysis, the presence of host-associated proteases (namely pepF, pepN, pepO, pepS, pepT) was confirmed in the two proteolytic active fractions (B8 and B10) but were not observed in the two fractions showing no proteolytic activity (B4 and B6).

**Figure 5.**
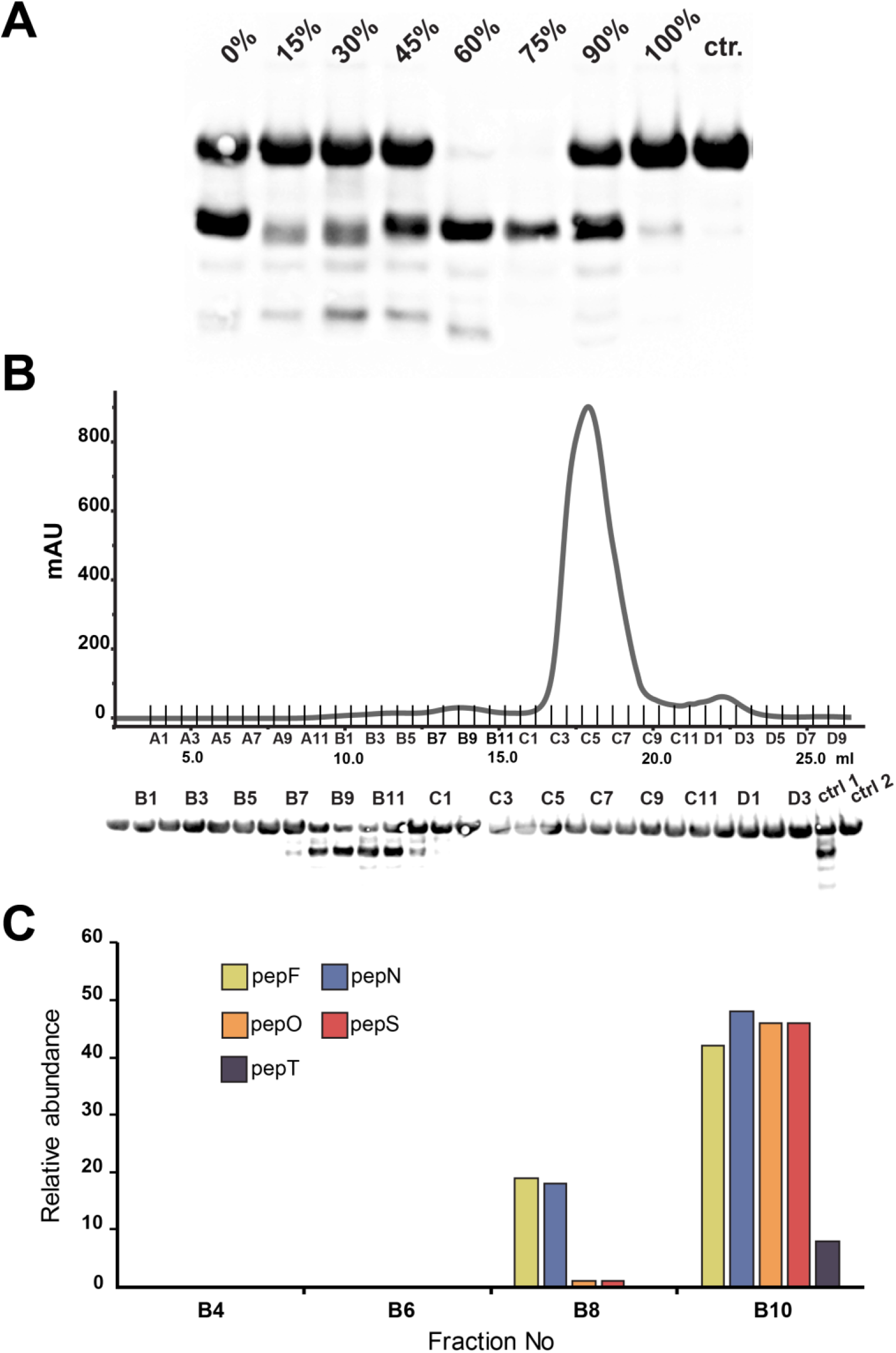
Identification of the proteases involved in the PlySK1249 cleavage. A) Proteins present in the total cell wall extract were serially precipitated using increasing concentrations of ammonium sulphate. Between each step, proteins were collected by centrifugation and then mixed with 40 μM of the LysM_CHAP construct and incubated overnight. Samples were then migrated on a NuPAGE 4-12% BisTris gel and degradation was confirmed by western blotting using Anti-6X His tag antibody. B) The 55% to 80% ammonium sulphate precipitation fraction was further separated using size exclusion chromatography. Fractions were tested for activity as described before. C) Relative abundances of the protein content of fractions B4, B6, B8 and B10 were analysed by MS-MS in order to observe an enrichment in protease related proteins. Enrichment was observed for the peptidases pepF, N, O, S, T. These were absent in the inactive B4 and B6 fractions but present in the active B8 and B10 fractions.

### Endolysin cleavage also occurred during *in vivo* prophage induction

As a proof of concept, we further tested whether endolysin specific cleavage products could also be generated during phage induction *in vivo*. Unfortunately, we could not induce the prophage carrying PlySK1249 from its natural *S. dysgaactiae* host. Therefore, we took advantage of *Streptococcus agalactiae* strain FSL S3-026, for which full genome is available (https://www.ncbi.nlm.nih.gov/nuccore/AEXT00000000.1) and which contains a single LambdaSa04-like prophage that is inducible with mitomycin C. This prophage carries an endolysin closely related to PlySK1249 (further named PlySK1249*; see Methods section and Supplementary Fig. 4A), including the same domain architecture and 80% aa homology with the PlySK1249 endolysin (Supplementary Fig. 5B).

Cultures of *S. agalactiae* FSL-S3 were either induced with mitomycin C or left uninduced as negative controls (Fig. 6A). Supernatants were then collected after 6h and loaded on an SDS-PAGE gel (Fig. 6B). The gel was split into six different fragments covering a range of molecular weights from 250 to 8 kDa and further analyzed by LC-MS for the presence of prophage proteins. A total of 11 prophage-related proteins were identified in the supernatant of the induced culture and, to a much lesser extent, in the non-induced fractions, suggesting basal induction of the prophage (Supplementary Table 3). A closer look at the peptides that were detected in the different gel fragments indicated that PlySK1249* was present in the supernatant in its original form, but also in specific cleavage products (Figure 6C). Peptides covering the total lengths of the enzyme were detected in band A (ranging from approximately 75 to 40 kDa) mainly in induced cultures and somewhat in non-induced samples. Peptides covering the Ami_LysM and Ami catalytic sites were also observed in the lower part of the gel at two different positions, indicating cleavage in the linkers on both sides of LysM. This fulfilled the concept that linker-domain specific cleavage of PlySK1249* endolysin did occur *in vivo* as observed with its PlySK1249 homologue *in vitro*. Finally, specific degradation was also confirmed when compared to the degradation patterns of the other phage proteins, for which only unspecific degradation was observed (Supplementary Fig. 5). These results further support the specificity of the endolysin cleavage sites.

**Figure 6.**
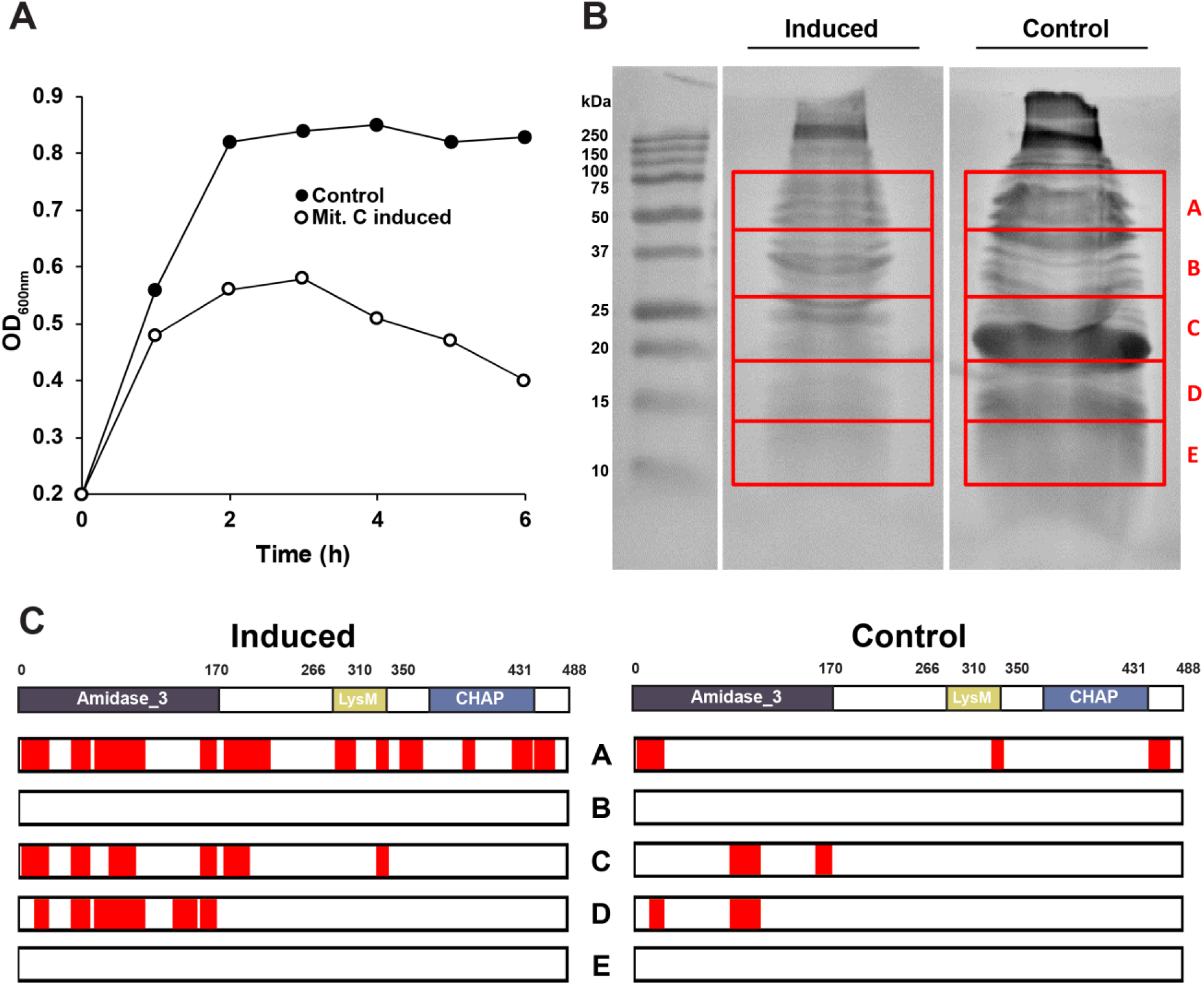
PlySK1249-like endolysin proteolysis in vivo during prophage induction in strain S. agalactiae FSL-S3. A) The LambdaSa04-like prophage inserted in the *S. agalactiae* FSL-S3 strain was induced by adding 1μl of mitomycin at an OD_600nm_ of 0.2 and lysis of the culture was followed for six hours (full circle non-induced, empty circle mitomycin C induced culture). B) The supernatant of the culture was then concentrated 5000x time and migrated onto a 15% SDS gel. A total of six bands were cut from the gel, covering a molecular weight range of 250 to 10 kDa. C) Extracted bands from the gel were analysed by LC-MS for the presence of the PlySK1249-like endolysin in both induced and un-induced cultures. Peptides detected that matched with the PlySK1249-likeamino acid sequences are highlighted in red.

### Possible implication of proteolytic cleavage for endolysin activity

The impact of proteolytic cleavage on the activity and diffusion of PlySK1249 was finally investigated (Fig 7). Incubation of PlySK1249 with trypsin for half an hour resulted in a reduced lytic activity on *S. dysgalactiae* SK1249 cells. Indeed, a turbidity decrease of only 40% was observed after 30 min compared to an 80% decrease after 10 minutes without trypsin (Fig. 7A).

**Figure 7.**
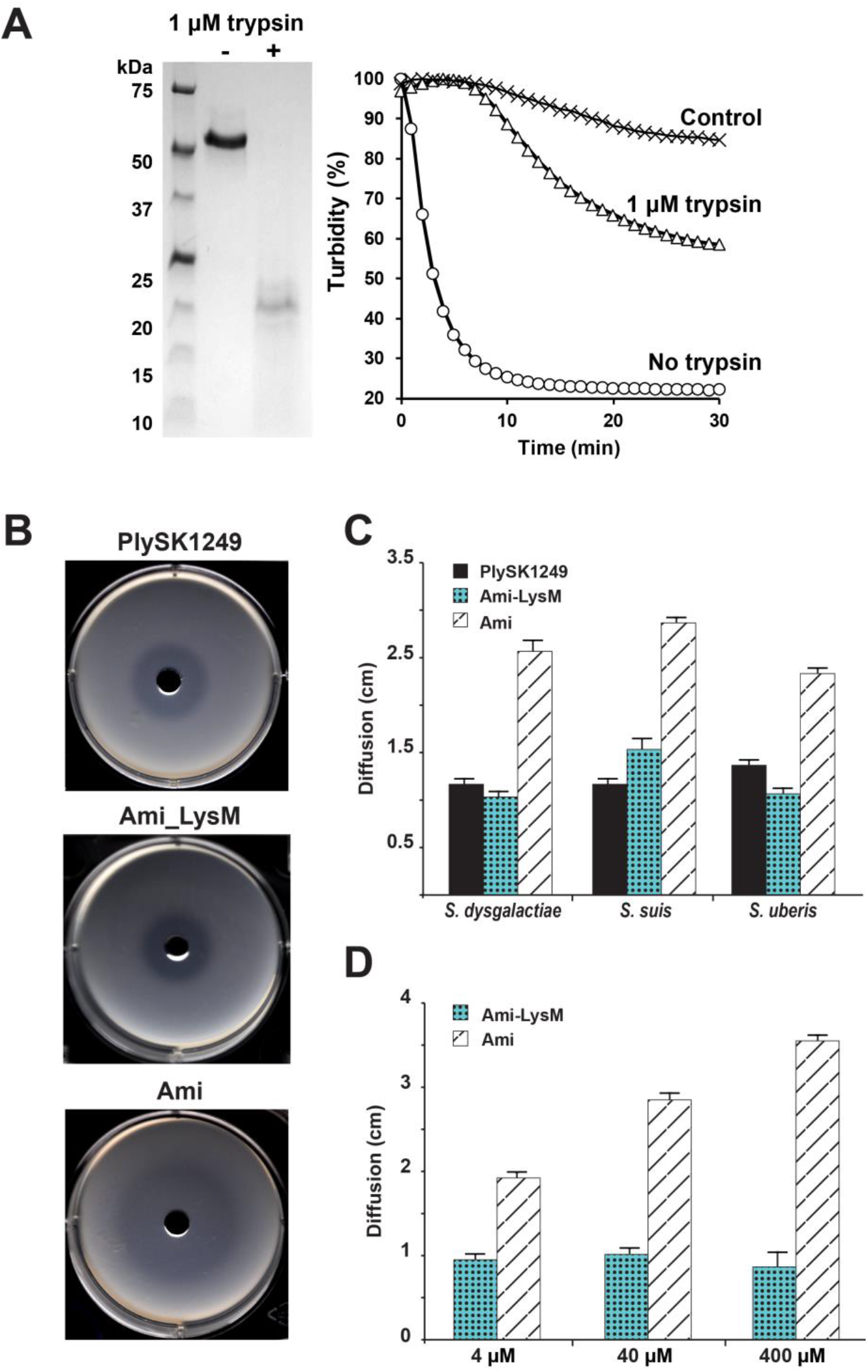
Effect of proteolytic cleavage on PlySK1249 lytic activity and impact of endolysin truncation on diffusion. A) *S. dysgalactiae* cells in the exponential growth phase were exposed to 3.5 μM of PlySK1249 or 3.5 μM of PlySK1249 pre-treated with 0.1 uM/ml of trypsin during 30 min. Decreases in bacterial cell turbidity was measured at 600 nm during 30 min. B) PlySK1249, Ami and Ami_LysM (40 μM) diffusion across a layer of soft agar containing heat inactivated *S. dysgalactiae* cells with formation of a lysis halo. C and D) Diffusion of the amidase domain was compared to the Ami_LysM construct and the native enzyme at a concentration of 40 μM and on different streptococcal species (C) or at different concentrations for the Ami and Ami_LysM constructs (D). Each experiment was repeated three times. Means ± standard deviations are shown.

Because proteolytic cleavage also resulted in the separation of the LysM binding domain from the catalytic sites, we decided to measure the diffusion of PlySK1249, Ami and Ami_LysM across a layer of soft agar containing heat inactivated *S. dysgalactiae* SK1249 cells. Due to the lytic activity of the amidase domain, it was possible to evaluate the diffusion of the different constructs by measuring the diameters of the clear halos produced by bacterial lysis (Fig. 7B). LysM clearly prevented diffusion of the Ami domain when tested against three different streptococci (average diameter decreased from ~2.5 cm for Ami compared to ~1 cm for Ami_LysM, Fig. 7D and 7E). The diffusion of PlySK1249 was similar to that of Ami_LysM, indicating that the phenomenon was independent to the protein molecular weight (Fig. 7D and 7E). In addition, diffusion of Ami was concentration-dependent, whereas LysM prevented diffusion over a >100-fold range of concentrations (Fig. 7F).

## Discussion

Endolysins of phages infecting Gram-positive bacteria usually harbor a two-domain structure with a CD at N-terminal and a CBD at the C-terminal end of the protein. Involvement of CDs and CBDs in cell lysis has been well documented and their complex regulation has been thoroughly investigated at the level of expression, translocation and activation (1,35,36). Recruitment of multiple (usually two) CDs has been identified as a potential evolutionary mechanism to acquire substrate specificity (37). However, different studies reported that multimodular endolysins may reunify both bacteriolytic and non-bacteriolytic catalytic domains, thus questioning the role of the non-bacteriolytic moieties in the endolysin function (7,16–19). The *S. dysgalactiae* endolysin PlySK1249 is another example of such bi-functional lysin with the association between a CHAP and an amidase domain (27). Here we aimed at understanding the individual and cooperative roles of the PlySK1249 functional domains, as well as their potential involvement in endolysin lysis regulation.

Using the parent enzyme and truncated constructs, we found that the three domains cooperated to increase the overall lytic activity of the enzyme. Although CHAP was not bacteriolytic, it did have a substantial cooperative effect (which was in fact synergistic since the activity of combined domains was greater than the sum of the activities of individual domains) when combined with Ami as in the context of native PlySK1249.

To better understand the contributions of Ami and CHAP in this cooperation, we further investigated the functions of the three domains and could demonstrate that the non-bacteriolytic CHAP domain had a de-chaining activity that also contributed to peptidoglycan solubilization. CHAP was a L-Ala-D-Ala endopeptidase that resolved complex polymers of stem-peptides to dimers and helped the Ami domain to digest peptidoglycan to completion. Moreover, resolving polymers to dimers rather than to monomers explained the lack of CHAP bacteriolytic activity, as dimeric stem peptides could keep glycan chains cross-linked and ensure a minimal cell wall skeleton. In contrast, complete digestion to monomers would have let the network of glycan chains fall apart, resulting in bacteriolysis.

The CHAP de-chaining activity is reminiscent of cell wall maturation by autolysins in *Bacillus subtilis*, which also act as de-chaining enzymes, as well as the Cse autolysin of *Streptococcus thermophilus*, which is composed of a LysM and a CHAP domain that promotes cell separation (30,31). Not unexpectedly, functionally common domains may serve different purposes in different backgrounds. While functional dissection of endolysin subdomains has been reported before, delving into the enzymatic contribution of each individual subdomains helped to reconcile the synergistic contribution of the non-bacteriolytic CHAP to the overall endolysin activity and wall solubilization. Although silent in terms of cell lysis, the CHAP domain is a genuine endopeptidase actively complementing the Ami domain, and was not an inactive evolutionary by-product on its way to being lost, as sometimes suggested (15,20).

This intramolecular domain cooperation could be modulated by the action of host cell wall associated proteases. We could observe specific proteolytic cleavage taking place in the linkers connecting both CDs to the central LysM CBD when PlySK1249 was co-incubated with cell wall associated proteases or during cell lysis by prophage induction. This potential regulatory mechanism could serve the purpose of dismantling domain proximity – and domain cooperation – therefore modulating the lytic activity of the enzyme. A similar phenomenon was previously observed for the main autolysins AtlA of *Enterococcus faecalis* and AcmA of *Lactococcus lactis*, which were also degraded by cell wall proteases (32–34). In the case of AcmA, proteolytic cleavage resulted in a reduced enzymatic and cell wall binding activity (32,33,38)

In summary, the presented observations provide a rationale for the multi-modular architecture of PlySK1249, which could possibly be extended to other multi-modular lysins. While both catalytic domains were observed to act coordinately to optimize bacterial lysis, the CBD is expected to delay diffusion until proteolytic inactivation of the lysin by host cell wall-associated peptidases is completed. This could also possibly protect neighboring bacteria and sibling prophages from deleterious lysis. Moreover, the multimodular structure of PlySK1249 might provide a way to channel several functions in one single protein as subsequent protein cleavage or maturation would free the modules to perform different functions, i.e. Ami for lysis and CHAP for de-chaining.

These observations have both theoretical and practical implications. From the theoretical point of view, it raises the question of coevolution between endolysin / autolysin and host bacteria. Endolysins and autolysins are likely to have common ancestors, and bacteria can regulate the activity of autolysins post-translationally (i.e. in the peptidoglycan) for the purpose of cell wall maturation. It now appears that the phage endolysin studied herein undergoes similar bacterial-dependent post-translational modification, highlighting the possible adaptation of phage endolysins toward the use of existing bacterial regulation mechanism for their own advantage.

From the practical point of view, it opens new perspectives on the ideal architecture of therapeutic antibacterial endolysins. Indeed, this is particularly relevant for the design of new chimeric endolysin, where domains that do not show lytic activity should not be necessarily dismissed. Indeed, when combined with appropriate partners, they could enhance the lytic properties of other CD domains. Moreover, the implication of the linkers should not be underestimated and further investigated in regards of enzyme stability during therapeutic use.

## Methods

### Cell culture and growth conditions

All bacterial strains used in this study are listed in Table 1. Gram-positive bacteria were grown at 37°C in Brain Heart Infusion (BHI, Becton Dickinson, New Jersey, USA) broth and plated on Mueller Hinton agar with 5% sheep blood (bioMérieux SA, Marcy l’Etoile, France). Broth cultures of streptococci and enterococci were grown without aeration. *Escherichia coli* were cultured at 37°C in Lysogeny Broth (LB) with agitation (220 rpm) or plated on LB Agar (LA). The following compounds (at final concentrations) were added to the media when necessary: kanamycin sulfate (30 μg/mL, chloramphenicol (25 μg/mL for LA plates and 50 μg/mL for LB) or isopropyl β-D-1-thiogalactopyranoside (IPTG) (0.4 mM). Culture stocks were prepared from cells in the exponential growth phase in 20% glycerol (vol/vol) and stored at −80°C. All chemicals were reagent grade, commercially available products.

### Cloning, expression and purification of the truncated versions of the PlySK1249 enzyme

The plasmid pPlySK1249^28a^ (27) was used as a template for PCR amplification of genes encoding for truncated forms of the PlySK1249 endolysin (Fig. 1A) using specific primer pairs listed in Table 1 (Microsynth AG, Balgach, Switzerland). PCR products were digested using restriction enzymes NcoI and XhoI (Promega, Madison, WI, USA) and ligated into pre-digested expression vector pET28a. Obtained plasmids (namely, pAmi^28a^, pAmi_LysM^28a^ and pLysM_CHAP^28a^, Supplementary Table 1), were transformed in One Shot BL21 (DE3) pLysS chemically competent *E. coli* cells (Life Technologies Europe B.V., Zug, Switzerland). Plasmid DNA was purified using the Qiagen Miniprep kit following the manufacturer’s recommendations and all constructs were validated by DNA sequencing using universal T7 primers (Supplementary Table 1). Following 0.4 mM IPTG induction for 18 hours, recombinant proteins were purified by affinity chromatography as described previously (27) and loaded on NuPAGE 4-12% BisTris gels (Invitrogen, Carlsbad, CA, USA) to confirm the correct molecular weights and assess their purity.

### Evaluation of the antibacterial efficacy of lysins

Efficacy of bacterial lysis was measured for the parent PlySK1249 endolysin and its various truncated constructs by following the decrease in OD_600nm_ of a bacterial suspension as previously described (27). In order to assess the potential interaction between Ami_LysM or Ami with LysM_CHAP, the truncated enzymes were mixed together at different molar ratios ranging from 1:1 to 1:100. *In vitro* time-kill assays were performed as described elsewhere (27).

### Light and electron microscopy

*S. dysgalactiae* cells were incubated with the various endolysin constructs at a concentration of 3.5 μM for 7 min for the PlySK1249 and Ami_LysM constructs, and for 1 h for the control and the CHAP domain. For light microscopy, cells were immobilized on 1% agarose pads and phase contrast microscopy images were taken with a Plan-Apochromat 100X/1.45 oil Ph3 objective on an AxioImager M1 microscope (Zeiss). For Transmission Electron Microscopy (TEM), cell suspensions were treated with glutaraldehyde (25% final concentration) for 1 h at room temperature (RT) and washed with phosphate-buffered saline (PBS). The same procedure was applied for metaperiodate (1% final concentration, 15 min incubation at RT) and osmium tetroxide plus hexacyanoferate (1% and 1.5%, respectively, 1 h incubation at RT). Cells were then centrifuged and the pellets were spun down in microcentrifuge tubes containing melted agar. After solidification of the agar, pellets were embedded in epon for ultra-thin sections that were prepared as described by Broskey et al. (39). Micrographs were taken with a transmission electron microscope FEI CM100 (FEI, Eindhoven, the Netherlands) at an acceleration voltage of 80 kV with a TVIPS TemCam-F416 digital camera (TVIPS GmbH, Gauting, Germany).

### Purification and digestion of peptidoglycan

Peptidoglycan of *S. dysgalactiae* SK1249 was purified as previously described (40). Briefly, 10 ml of an overnight culture was added to 1 L of fresh BHI medium. The cells were grown until an OD_600nm_ of 0.4-0.5 and then quickly cooled in an ice bath for 5-10 min. The culture was centrifuged at 4°C and resuspended in PBS to reach a total volume of 40 ml. The bacterial suspension was poured dropwise into 40 ml of boiling sodium dodecyl sulphate (SDS) (8%) and boiled with agitation for 15 min to inactivate intrinsic autolytic enzymes. The cells were centrifuged at 20°C to avoid SDS precipitation, washed twice with NaCl 1M, and five times with dH_2_O. After the final washing step, the bacterial pellet was resuspended in 2 mL dH_2_O and stored at −20°C overnight. The following day, the cells were broken using a FastPrep homogenizer (Thermo Savant FastPrep FP120 Homogenizer) during three bursts of 45 sec at 6.5 m/sec with a 5 min cooling step between each burst. The supernatant was transferred into a tube and centrifuged at 13’000 rpm for 5 min at 4°C and then again for 20 min at 4°C. The pellet was resuspended in 3 mL Tris (0.1 M), 0.3 mL NaN_3_ (0.5%), pH 7.5. After the addition of 0.3 mL MgSO_4_ (200 mM), 60 μl DNAse (0.5 mg/mL) and 60 μl RNase (2.5 mg/mL), the mixture was incubated for 2 h at 37°C with agitation, and then overnight after 0.3 mL CaCl_2_ (100 mM) and 0.3 mL trypsin (100 μg/ml) were added. For the final steps, SDS (1% final concentration) was added to the preparation, which was then heated for 15 min at 75°C to extract digested peptides. After an additional centrifugation for 20 min at 20°C and 13’000 rpm, the pellet was washed once with dH_2_O, resuspended in 20 mL LiCl (8 M) and incubated at 37°C with agitation for 15 min. The mixture was again centrifuged at 13’000 rpm, 4°C for 20 min. Pellet was resuspended in 20 mL EDTA (0.1 M) to remove material bound by ionic interactions. Lastly, the bacterial peptidoglycan was centrifuged, washed twice with 2 mL dH_2_O and resuspended in 2 mL dH_2_O. Aliquots of 1 mL were transferred into pre-weighed Eppendorf tubes and dried overnight by rotary evaporation (UniEquip UNIVAPO 150 ECH). The dried bacterial peptidoglycan was then resuspended to reach a concentration of 10 mg/mL. For enzymatic digestion, 100 μl of the extracted peptidoglycan was mixed with 900 μl of the previously purified endolysin constructs at 3 μM. The mixture was incubated overnight at 37°C with agitation. The following day, the solution was heated for 3 min at 100°C to inactivate the endolysin, then centrifuged at 13,000 rpm, RT for 10 min, and the supernatant containing the solubilized walls was transferred into a new tube.

### Biochemical analysis of PlySK1249 hydrolysis sites in the peptidoglycan

Peptidoglycan extraction was performed as described above. Reducing sugar analysis was conducted using a modified Park-Johnson assay, and free amino acids were quantified with a modified Ghuysen procedure with 1-fluoro-2.4-dinitrobenzene, as described by Schmelcher et al. (41). Undigested peptidoglycan was used as a blank and glucose or L-Alanine calibration curves were used for calculations. All experiments were performed in triplicate.

### Analysis of the digested peptidoglycan using Reverse Phase - High Pressure Liquid Chromatography (RP-HPLC)

The digested peptidoglycan was kept at −80°C for 5 min and then dried overnight by rotary evaporation. After the addition of 0.5 mL of acetone, the tube was sonicated to re-suspend the pellet and another 1 mL of acetone was added to remove any contaminating endotoxins. The tube was left at RT for 30 min and then centrifuged at 13’000rpm, RT for 10 min. The supernatant was dried by rotary evaporation for 10 min. 200 μL dH_2_O was added before sonication. Samples was stored at −80°C for 5 min and dried by rotary evaporation for 2 h. Pellet was resuspended at 2 mg/mL in 25% 2-propanol, 25% acetonitrile, 50% dH_2_O, 10% TFA to separate the glycan chains from the peptides by differential precipitation. The tube was vortexed thoroughly, sonicated and placed on ice for 15 min. After centrifugation at 13’000 rpm for 15 min at 4°C, the supernatant containing the stem peptides and peptide bridges, was transferred into a new tube and stored at −80°C until further analysis. Before analysis, the frozen peptides were dried by overnight rotary evaporation and resuspended in 500 μL dH_2_O. To separate the peptides, an HPLC system (Hitachi Instruments, Ichige, Hitachinaka, Japan) consisting of an L-7200 autosampler, an L-7100 gradient pump and an L-7400 UV detector was used as described previously (40). Aliquots of 100 μL were injected into a C18 reverse phase column (SuperPac Sephasil C18, 5 μm, 4 × 250-mm column, Amersham Pharmacia Biotech) maintained at 25°C using a pelcooler (Lab-Source, Reinach, Switzerland). A linear gradient of 0-15% acetonitrile in 0.1% trifluoroacetic acid at a flow rate of 0.5 mL/min over 100 min allowed separation of the peptides, which were detected at 210 nm. Collected data were analyzed with the D-7000 HPLC System Manager program (Hitachi).

### Liquid Chromatography coupled with Mass Spectrometry (LC-MS) analyses of the digested peptidoglycan

Peptidoglycan extraction was performed as described above. At the end of the digestion, samples were boiled at 100°C to stop the enzymatic reaction. They were filtered through a 5’000 MWCO cut-off column (Vivaspin 500, Sigma, St. Louis, MO) and desalted using a C18 cartridge (Thermo Fisher, Bremen, Germany). Peptidoglycan fragments were eluted with an 80% MeCN, 0.1% TFA solution and dried by rotary evaporation, before being dissolved in loading buffer (2% MeCN, 0.1% TFA). Samples were injected on a Dionex RSLC 3000 nanoHPLC system (Dionex, Sunnyvale, CA, USA) interfaced via a nanospray source with a high-resolution mass spectrometer QExactive Plus (Thermo Fisher, Bremen, Germany). Peptidoglycan fragments were loaded onto a trapping microcolumn Acclaim PepMap100 C18 (20 mm x 100 μm ID, 5 μm, Dionex) before being separated on a C18 reversed-phase analytical nanocolumn at a flow rate of 0.25 μL/min. A Q-Exactive Plus instrument was interfaced with an Easy Spray C18 PepMap nanocolumn (50 cm x 75 μm ID, 2 μm, 100Å, Dionex) using a gradient programmed to run over 37 min from 4 to 76% acetonitrile in 0.1% formic acid. Full mass spectrometry survey scans were performed at 70,000 resolution. In data-dependent acquisition controlled by Xcalibur software 3.1 (ThermoFisher), the ten most intense multiply charged precursor ions detected in the full MS survey scan (250-2000 m/z window) were selected for higher energy collision-induced dissociation (HCD, normalized collision energy = 27%) and for analysis in the Orbitrap at 35,000 resolution. The window for precursor isolation was of 1.5 m/z units around the precursor and selected fragments were excluded from further analysis for 10 s. MS raw files were processed with PEAKS software (version 8.0, Bioinformatics Solutions Inc., Waterloo, Canada) and used for peptide identification by *de novo* sequencing and a database search. For the database search, a set of *S. dysgalactiae* (subspecies *equisimilis*) SK1249 proteome sequences was downloaded from the UniProt database (February 2016 version, 2,309 sequences) and used to identify sample contamination by proteins from *S. dysgalactiae*. At the same time, *de novo* sequencing was used to identify peptidoglycan peptides. The database search and *de novo* sequencing parameters were as follows. No enzyme was used as the enzyme definition, N-terminal acetylation of protein and oxidation of methionine were used as variable modifications, and a parent ion tolerance of 10 ppm and a fragment ion mass tolerance of 0.02 Da were used.

### Proteolytic degradation of the PlySK1249 endolysin in the presence of cell-wall protein extracts and trypsin

Cell wall protein extracts from the strain *S. dysgalactiae* SK1249 were prepared, as described elsewhere (42). PlySK1249 and its truncated versions (40 μM) were digested overnight at 37°C by mixing them with half volume of the extract. Additional endolysin samples were also digested by adding trypsin at 1 μM for different incubation times. Digestions were conducted at 37°, 200 rpm agitation. When necessary, purified peptidoglycan was added to the reaction at a final concentration of 100 μg/mL. The reaction was stopped by incubating the mix with loading buffer and beta-mercaptoethanol for 15 min at 80°C. The reaction products were loaded on a NuPAGE 4-12% BisTris gel. Bands of interest were cut from the gel and the amino acid sequences were determined by nanoLC-MS/MS at the protein analysis facility at the University of Lausanne (PAF, Center for Integrative Genomics, University of Lausanne, Switzerland). Western blotting was performed as described previously (43) and blots were incubated for 1 h with a 1:1,000 dilution of anti 6X-His Tag rabbit antibodies (Thermo Fisher Scientific, Massachusetts, USA) followed by a 1 h incubation with a 1:3,000 dilution of goat anti-rabbit IgG coupled to HRP secondary antibodies (Thermo Fisher Scientific). Bands were detected by chemiluminescence using ECL Western blotting detection reagent (Amersham Bioscience. Piscataway, NJ). For fractionation of the cell wall protein extract, ammonium sulfate was used at multiple concentrations ranging from 15% to 100%. For each fractionation step, finely ground ammonium sulfate was added directly to 10 mL of cell wall extract and agitated for 15 min at 4°C. The protein fractions were collected by centrifugation at 4000 rpm for 30 min. Pellets were resuspended in 100 μL PBS and mixed with 100 μL of the LysM_CHAP construct for overnight incubation. Western blotting was used to assess proteolyzed fragments in the different fractions, as described above. A second fractionation step was performed with 55% to 80% ammonium sulfate fractions loaded on a PBS-equilibrated size exclusion column (Superdex® 200 Increase 10/300 GL, GE Healthcare, Little Chalfont, United Kingdom). The protein content was eluted in 0.5 mL fractions with one column volume of PBS. Eluted fractions were tested for proteolytic activity using the LysM_CHAP construct, as described above. Four fractions were further analyzed by nanoLC-MS/MS at the protein analysis facility at the University of Lausanne (PAF, Center for Integrative Genomics, University of Lausanne, Switzerland).

### Endolysin proteolysis during *in vivo* prophage induction

For prophage induction, *S. dysgalactiae* strain FSL-S3-026 was grown at 37° in a synthetic CDEN medium (Amimed, Switzerland) to an OD_600nm_ of 0.2. Mitomycin was added at a final concentration of 1 μg/mL to induce prophage excision. After a 6 h induction, the culture was centrifuged, filtered at 0.22 μm and concentrated to 1 mL using a Vivaspin turbo ultrafiltration units (10 kDa MWCO, Sartorius AG, Goettingen, Germany). Samples were further concentrated to 50 μL after five successive washes using Tris 50 mM, pH 7.5 before being run on a 15% polyacrylamide SDS-PAGE gel. A total of six bands covering molecular weights ranging from 8 to 75 kDa were finally cut from the gel and peptides contained in each band were analyzed by nanoLC-MS/MS, as described above.

### Endolysin diffusion assay

This assay was designed to assess the implication of the CBD in the diffusion of the endolysins through a layer of PBS soft-agar inoculated with bacteria. An overnight culture of *S. dysgalactiae* SK1249 was washed in PBS and resuspended in 0.25 vol of PBS. Granulated agar (7.5 g/L) was added to the cell suspension and autoclaved for 15 min at 120°C. One mL/well of solution was poured into 6-well plates and stored at 4°C until further use. 4 mm diameter wells were punched in the center of the agar surfaces and 10 μL aliquots of the enzyme at different concentrations were added into the pits. Diffusion was assessed by measuring the diameter of the lysis halo that formed after overnight incubation at 37°C.

## Acknowledgments

The authors would like to thank Paul Majcherczyk for his outstanding technical help with the HPLC analysis, Partrice Waridel for running all the LC-MS analyses and Michael Taschner for helping with the size exclusion chromatography experiments. We are grateful to Sara Mitri and Harald Brüssow for fruitful discussions, comments and for reviewing the manuscript.

## Disclosure statement

The authors declare that they have no conflicts of interest with the contents of this article.

## Funding

This work was supported by a nonrestricted grant from the Foundation for Advancement in Medical Microbiology and Infectious Diseases (FAMMID) (P.M.). This work was also supported by the Swiss National Science Foundation (SNSF) under grant P400PB_191059 (F.O).

